# Phenopolis: an open platform for harmonization and analysis of sequencing and phenotype data

**DOI:** 10.1101/084582

**Authors:** Nikolas Pontikos, Jing Yu, Fiona Blanco-Kelly, Tom Vulliamy, Tsz Lun Wong, Cian Murphy, Valentina Cipriani, Alessia Fiorentino, Gavin Arno, Daniel Greene, Julius OB Jacobsen, Tristan Clark, David S Gregory, Andrea Nemeth, Stephanie Halford, Susan Downes, Graeme C Black, Andrew R Webster, Alison Hardcastle, Vincent Plagnol

**Author notes:** Contributed equally to the work. Centre for Ophthalmology & Vision Sciences, Institute of Human Development, Faculty of Medical and Human Sciences, University of Manchester Manchester, UK.

## Abstract

**Summary:** Phenopolis is an open-source web server which provides an intuitive interface to genetic and phenotypic databases. It integrates analysis tools which include variant filtering and gene prioritisation based on phenotype. The Phenopolis platform will accelerate clinical diagnosis, gene discovery and encourage wider adoption of the Human Phenotype Ontology in the study of rare disease.

**Availability and Implementation:** A demo of the website is available at http://phenopolis.github.io (username: demo, password: demo123). If you wish to install a local copy, souce code and installation instruction are available at https://github.com/pontikos/phenopolis. The software is implemented using Python, MongoDB, HTML/Javascript and various bash shell scripts.

**Supplementary information:** http://phenopolis.github.io

## INTRODUCTION

The molecular diagnosis of rare genetic diseases requires detailed clinical phenotypes and processing of large amounts of genetic data. This motivates large-scale collaborations between clinicians, geneticists and bioinformaticians across multiple sites where patient data are pooled together to boost the chances of solving rare cases, and validating novel genes. For example to solve retinal dystrophies, the UK Inherited Retinal Dystrophy Consortium (UK-IRDC) has set up a collaboration between London, Manchester, Oxford and Leeds.

A complication of multi-site collaborations is that discrepancies in phenotype definitions and interpretation of genetic variants can lead to different molecular diagnosis (Yen et al. 2016). A solution to reduce the variability introduced by different sequencing analysis pipelines is to analyse the sequence data centrally and store the annotated variants in a large normalised database. On the clinical side, phenotype harmonisation is improved by using nomenclatures such as the Human Phenotype Ontology (HPO) (P. N. Robinson et al. 2008) to translate specific clinical features into a standardised, computer interpretable format.

We have integrated these approaches into Phenopolis, an interactive website built on genetic and phenotypic databases. With the help of HPO-encoded phenotypes, Phenopolis is able to prioritise causative genes using different sources of evidence, such as published disease gene associations from the Online Mendelian Inheritance in Man (Hamosh et al. 2005), abstract relevance from Pubmed publications, as well as model organism phenotype ontology analysis using the Exomiser (P. Robinson et al. 2013), (Bone et al. 2016). Additionally, Phenopolis uncovers gene phenotype relationships within the stored patient data through variant filtering and statistical enrichment of HPO terms. The online version available at http://phenopolis.github.io, includes four example patients with inherited retinal dystrophies, to illustrate our methods.

### Clinical data collection

The collection of clinical phenotype data was done retrospectively from patient records and entered using the Phenotips platform (Girdea et al. 2013), which provides an interface for translating detailed clinical phenotypes into HPO terms. Various patient diagnoses were translated to their closest match using HPO terminology. This included mode of inheritance and modifiers such as age of onset and laterality when available.

### Genetic data collection

Our internal exome database, UCLex, currently comprises 4,449 patients which were collected from various research groups since 2012. Four patients solved with genetic mutations in *DRAM2* (El-Asrag et al. 2015) and *TTLL5* (El-Asrag et al. 2015) are made available on the demo account.

### Analysis of genetic data

The short read sequence data was aligned using novoalign, and variants and indels were then called according to GATK best practices (joint variant calling followed by variant quality score recalibration) (McKenna et al. 2010). The variants were then annotated using the Variant Effect Predictor (McLaren et al. 2016), output to JSON format, post processed by a Python script and loaded into a Mongo database.

### Website implementation

The Phenopolis website was implemented using the Python Flask web framework by extending the ExAC code base **[1]**. Javascript was used for visualisations (mostly using D3.js) and to make the website interactive (**Figure A**).

The website provides five main entry points:

- The home page: summary statistics of genetic and phenotypic data, as well as auto-completing search bar to search by phenotype, gene name or patient id.
- The all patients page: summary data of all patients and their candidate genes for which the user has access permission.
- The patient page: the patient phenotypes and a table of filtered variants per patient prioritised based on gene. The causal variants are expected to be in this list, ranked at the top of the table.
- The gene page: the variants and the patients in which they occur, as well as the gene-HPO analysis.
- The phenotype page: a prioritised list of genes per phenotype, based on known association and gene enrichment analysis.

## Applications

### Clinical application: gene prioritisation by patient

Given a list of genetic variants and the phenotype of a patient, the first task towards a molecular diagnosis is to prioritise potentially causative genes.

For each case, variants are first filtered based on user-defined thresholds:

- Allele count less than 5 in our internal database and in ExAC (M Lek 2016)
- Kaviar frequency less than 0.05
- Exclude non-exonic variants or variants on non-coding transcripts. Splicing variants are kept.

Next, gene panels from the gene to HPO/OMIM mapping **[2]**, available on the HPO website, and more specialised gene panels, such as Retnet **[3]** for retinal genes, are used to highlight candidate genes which match the phenotypic description and inheritance pattern. We have solved and validated several cases using this approach. We have also developed a Venn diagram visualisation to highlight genes which are associated to more than one phenotype (**Figure B**). We also provide a filterable variant table in which genes are ranked based on their Exomiser, Pubmed or Phenogenon gene scores (Supplementary).

We have successfully prioritized the *GNB3* gene in retinal dystrophy using the Pubmed relevance score that was able to leverage the evidence from relevant chicken model data (Arno et al. 2016).

### Research application: HPO signature per gene

Given a sufficiently large and phenotypically diverse collection of cases, gene to phenotype patterns start emerging. In order to assign phenotype associations per genes based on our patient database, we have developed a gene-based HPO enrichment and visualisation tool, Phenogenon (**Figure C**). We have also integrated the existing SimReg tool which suggests a characteristic phenotype per gene (Greene et al. 2016). Both methods work on a filtered list of variants and are explained in detail in the Supplementary section.

### Research application: genes per HPO term

Individuals with the specified HPO term and their solved gene are listed on this page. We retrieve the list of known disease genes from the gene-HPO/OMIM mapping **[2]** and we score these genes with Phenogenon to assess their support in our dataset. Furthermore, we rank all genes according to their Phenogenon score for this HPO term to enable gene discovery in our dataset.

## DISCUSSION

There are currently several closed-source commercial online alternatives that provide variant filtering and prioritisation, for example Saphetor **[4]**, Congenica **[5]** and Omicia **[6]**. However their tariffs are often expensive and they are not readily extensible. There are also open-source alternatives such as Seqr **[7]** and Gemini (Paila et al. 2013) but neither provide full integration with HPO terms yet.

As it stands, Phenopolis is an ideal platform for studying the pleiotropy of genes: how variation in different parts of the same gene could lead to different seemingly unrelated phenotypes (Supplementary).

In the next iteration of our software, we will be integrating tissue expression databases, allowing for genes and transcripts to be prioritised by cell type when the disease affects a specific tissue type. Furthermore, we are working on including CNV data, inferred from our exomes using ExomeDepth (Plagnol et al. 2012), as these can also be gene disrupting mutations.

We also plan on interfacing with the Genomics England GenePanel app to retrieve relevant genes and contribute novel disease genes.

Collection of phenotypes and prioritisation of genes can help elucidate which features are informative for a particular gene and warrant close inspection in clinic. The systematic chronological ordering of patient features obtained from clinical history can be very informative in discerning between conditions which might appear similar, for example rod-cone and cone-rod dystrophy. Currently, a limitation to obtaining detailed phenotypes for our retrospective cases is the manual input of HPO terms and we are investigating data-mining of health records to pull data efficiently.

Given the utility of this software within the UK-IRDC, we hope it will be used by other groups collaborating on the genetics of rare diseases.

## Acknowledgments

We wish to acknowledge all patients that contributed their DNA to this project. We also thank and acknowledge TS and DSG, for the Computer Science High Performance Compute which providing us with the computing infrastructure on which to analyse our data and host our webserver. We also wish to thank Michael Morgan for his insightful advice on Phenogenon.

NP and JY wrote the manuscript and it was critically reviewed by VC, ARW and DG. CM did the population PCA plot on the home page and performed the per phenotype gene burden test. DG carried out the SimReg analysis. JJ and VC helped with the Exomiser analysis. FBK, TV, VC, JY and GA provided the clinical patient data in HPO format. TC and DSG provided the computing infrastructure and the server configuration.

## URLs

[1] ExAC code base: https://github.com/konradjk/exac_browser

[2] Gene-HPO download: http://compbio.charite.de/jenkins/job/hpo.annotations.monthly/lastStableBuild/artifact/annotation

[3] Retnet: https://sph.uth.edu/Retnet

[4] Saphetor: www.saphetor.com

[5] Congenica: www.congenica.com

[6] Omicia: www.omicia.com

[7] Seqr: https://seqr.broadinstitute.org/

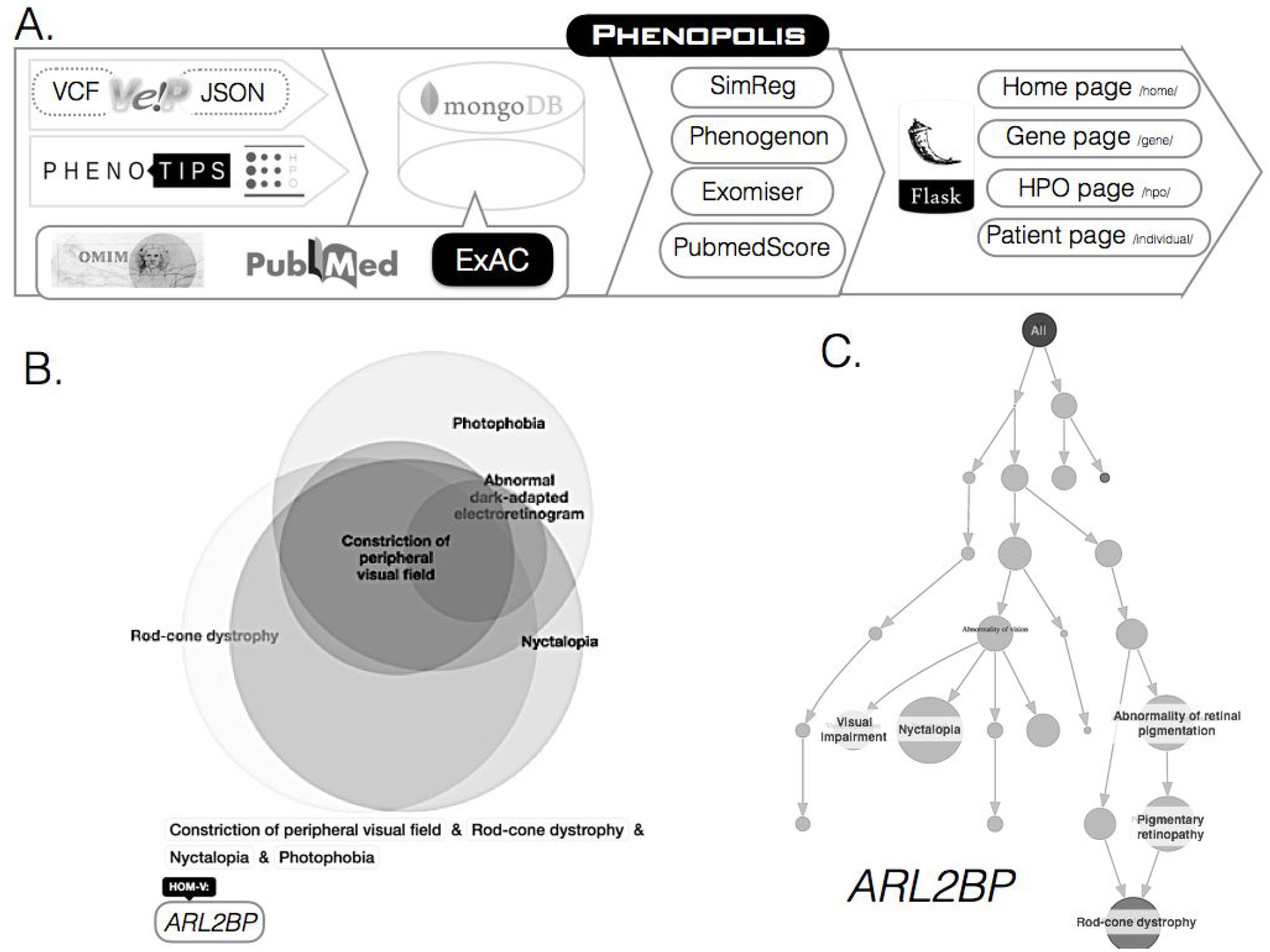
A. **Overview of the pipeline.** Phenotypic information is entered using Phenotips. The Variant Call Format files are annotated by the Variant Effect Predictor and translated to JSON for convenient import to MongoDB. OMIM, Pubmed and ExAC data are also imported into the Mongo database, on which we run the PubmedScore, Exomiser, SimReg and Phenogenon to score the genes. A Python Flask server is used as the front-end to display the four entry points to the website. **B. Venn diagram visualisation.** Genes at the intersection of multiple HPO-gene sets are highlighted. **C. The Phenogenon visualisation.** The size of the circles is inversely proportional to the p-value. Clicking on the nodes brings up information about the individuals and variants.

## Supplementary

### HPO Venn Diagram

A Venn diagram visualisation of the HPO to gene mapping is rendered with Javascript D3. Only genes for which there are filtered variants in the patient are displayed by default. It is also possible to view all genes by selecting “Toggle All”. Since the computation of the Venn diagram scales exponentially with the number of HPO terms, we currently limit to the five most specific terms.

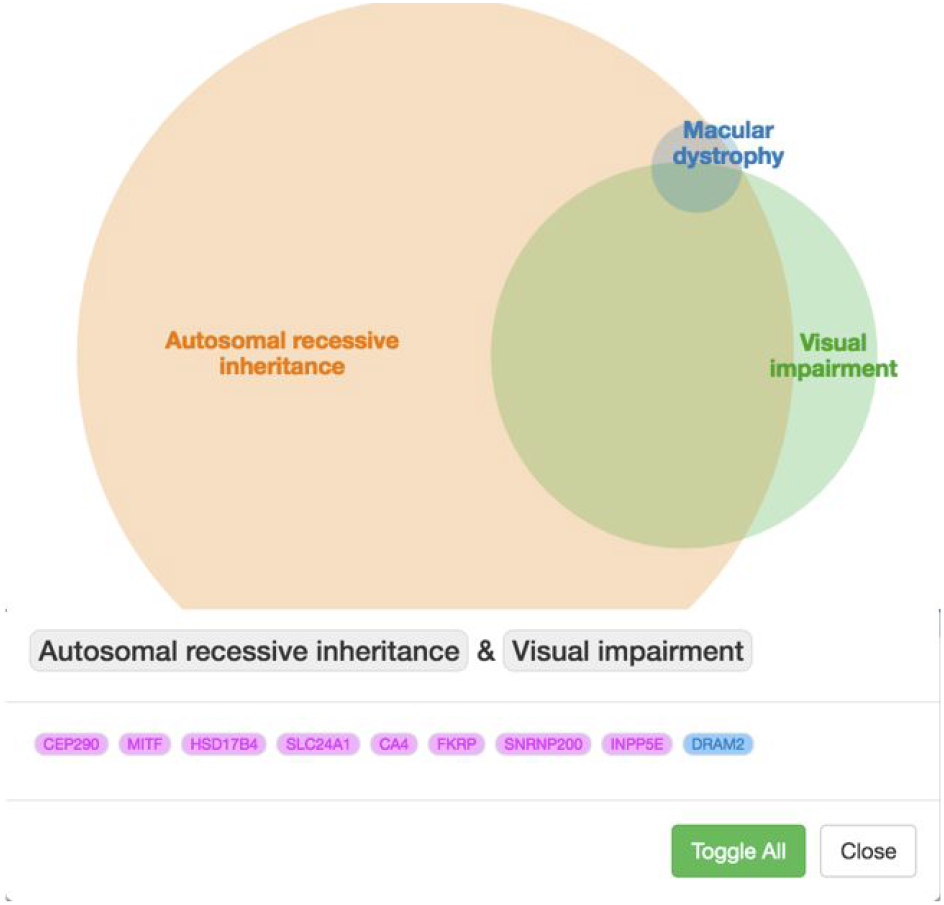

### PubmedScore

Genes are prioritised by PubmedScore calculated using literature support. PubMed publications are mined for interesting phenotypes given relevant keywords. Users may also supply a list of genes of interest that are known to be associated with the phenotype.

PubmedScore will assign at least a score of 1 to the genes on the given list. This will help to keep those genes on top of the result table if the user choose to sort it by the Pubmed Relevance Score.

Define an indicator function *B_known_*, given a gene (*g*):

*B_known_*(*g*) = 1: if the gene is on a given list of genes known to be associated with the phenotype

= 0: otherwise

The *pubmedscore*(*g*) is calculated by counting the appearance of each keyword (user defined) in every returned pubmed paper’s title and abstract, and taking the sum.

The overall Pubmed Relevance (*Sp*) score of a given gene (*g*) is calculated as:

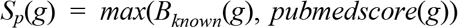

One can also insert an additional function adapted to problems at hand. Taking patients with retinal dystrophy as an example: Retnet (https://sph.uth.edu/Retnet/) has detailed information for retinal dystrophy associated genes, such as if a gene may cause dominant or recessive inheritance mode of a disease. This can then be used to help calculate *S_p_*. Given:

*B_ret_*(*g*) = 1: if the gene is reported on RetNet

= 0: otherwise

*B_retMode_*(*g*) = 1: if the gene is registered on RetNet and changes on the gene is described therein as possibly causing Dominant or X-linked or M-linked diseases

= 0: otherwise

*B_searchMode_*(*g*) = 1: if it is searched in the context of recessive inheritance mode

= 0: otherwise

Then

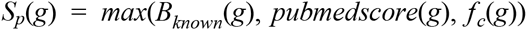

where

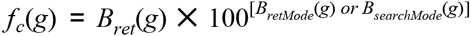

The function is then guaranteed to rank the known retinal dystrophy associated genes on top of the table that matches the inheritance mode.

Take gene DRAM2 as an example (can be found as candidate gene for two of the demo patients): Pubmed search with keywords ‘blindness macula macular pigmentosa retina retinal retinitis stargardt’ returns three publications (pubmed IDs: 27518550, 26720460, 25983245). The app then counted the appearance of the keywords in the titles and abstracts of the three publications, which equaled to 16. The *pubmedscore*(*DRAM*2) was then assigned as 16. However, since *DRAM2* was registered on RETNET as a recessive gene, and this patient has a relevant homozygous change on *DRAM2*, therefore

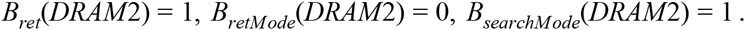

One can derive that

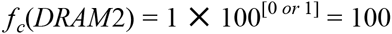

And hence the Pubmed Relevance score of *DRAM2* is

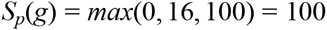

### Exomiser

When mouse, drosophila or other model organism phenotype ontologies are available, Exomiser can help find orthologous genes, by doing cross-species phenotype comparisons using the OWLSim algorithm (https://github.com/owlcollab/owltools). This works particularly well for embryonic development genes such as *PAX6* (P Robinson 2013, D Smedley 2015), whose function are conserved across species.

Resulting variants were processed with Exomiser (Robinson et al. 2014), a variant prioritisation software that annotates, filters and prioritises likely causative variants starting from a VCF file and a set of phenotypes encoded using the Human Phenotype Ontology (HPO) (Robinson et al. 2008). The functional annotation of variants was handled by Jannovar (Jager et al. 2014) that is embedded within Exomiser and uses UCSC KnownGene transcript definitions and hg19 genomic coordinates. Variants listed in the VCF files were first filtered according to user-defined criteria on rarity (minor allele frequency (MAF) ≤ 0.001 in either the publicly available 1000 Genomes Project Phase 1 component of dbSNP (1000G), the NHLBI GO Exome Sequencing Project and the Exome Aggregation Consortium datasets (ExAC)), autosomal dominant mode of inheritance (heterozygote) and quality (Phred Quality score Q > 30). The filtered variants were then ranked on the basis of their deleteriousness (predicted pathogenicity data as extracted from the dbNSFP resource (Liu et al. 2011)), and phenotypic relevance: that is how closely the given HPO-encoded phenotype matches the known phenotype of disease genes from human (Amberger et al. 2011; Maiella et al. 2013), mouse (Bult et al. 2013; Smith et al. 2005) and zebrafish (Van Slyke et al. 2014) model data (cross-species phenotype comparisons performed by PhenoDigm tool (Smedley et al. 2013).

### Phenogenon

Unrelated patients with observed HPO terms are included in the analysis, the number of which is denoted as Pata. The number of patients affected by a given HPO term *h* is denoted as Path. The number of patients who have a specific genotype in a gene *g* is denoted as Patg (this may be a single variant or more than one for compound hets). The variants are filtered by ExAC allele frequency and CADD phred score (M Kircher 2014) according to user specified thresholds so that only rare, predicted, damaging variants are included (frameshift variants which do not have a CADD score are assigned a default score of 50). The number of patients having both HPO term *h* and filtered variants in *g* is denoted as Patgh. Therefore one can construct a 2 × 2 table as follows, per gene *g* and HPO term *h*:

| | Number of patients **not** affected by $h$ | Number of patients affected by $h$ |
| --- | --- | --- |
| Number of patients **without** variant in $g$ | $Pat_a - Pat_h - Pat_g + Pat_{gh}$ | $Pat_h - Pat_{gh}$ |
| Number of patients with variant in $g$ | $Pat_g - Pat_{gh}$ | $Pat_{gh}$ |

Fisher’s Exact test is used to test the non-independence between *g* and *h*, and the phi correlation coefficient is used to quantify the correlation between *g* and *h*. Fisher’s Exact test produces two p values: right-tail and left-tail p values. A small right-tail p value indicates a positive correlation between *g* and *h*. The sign and magnitude of phi also determines the type of correlation. A strong positive phi indicates that rare damaging variants in the gene appear more often than expected by chance in patients with that HPO term. This suggests the disrupted gene may cause the phenotype. A negative phi would indicate a gene has less rare damaging variants than expected by chance, which might suggest a phenotype is driven by a more functional gene.

Obviously this approach is currently susceptible to sequencing bias. For instance, if all individuals which share a particular HPO term have improved coverage for a given gene then there will be an enrichment of rare variants in that gene. We are currently working on extending the method to account for this.

This method is applied to the gene with two possible inheritance modes, dominant where patient has to have at least one qualified variant, and recessive where the patient has to have at least two qualified variants. The significance of the top p-values can also help infer whether a gene is more likely to cause dominant or recessive mode of the phenotype.

The HPO tree view also nicely illustrates syndromic genes which cause a range of phenotypes, for example *USH2A*:

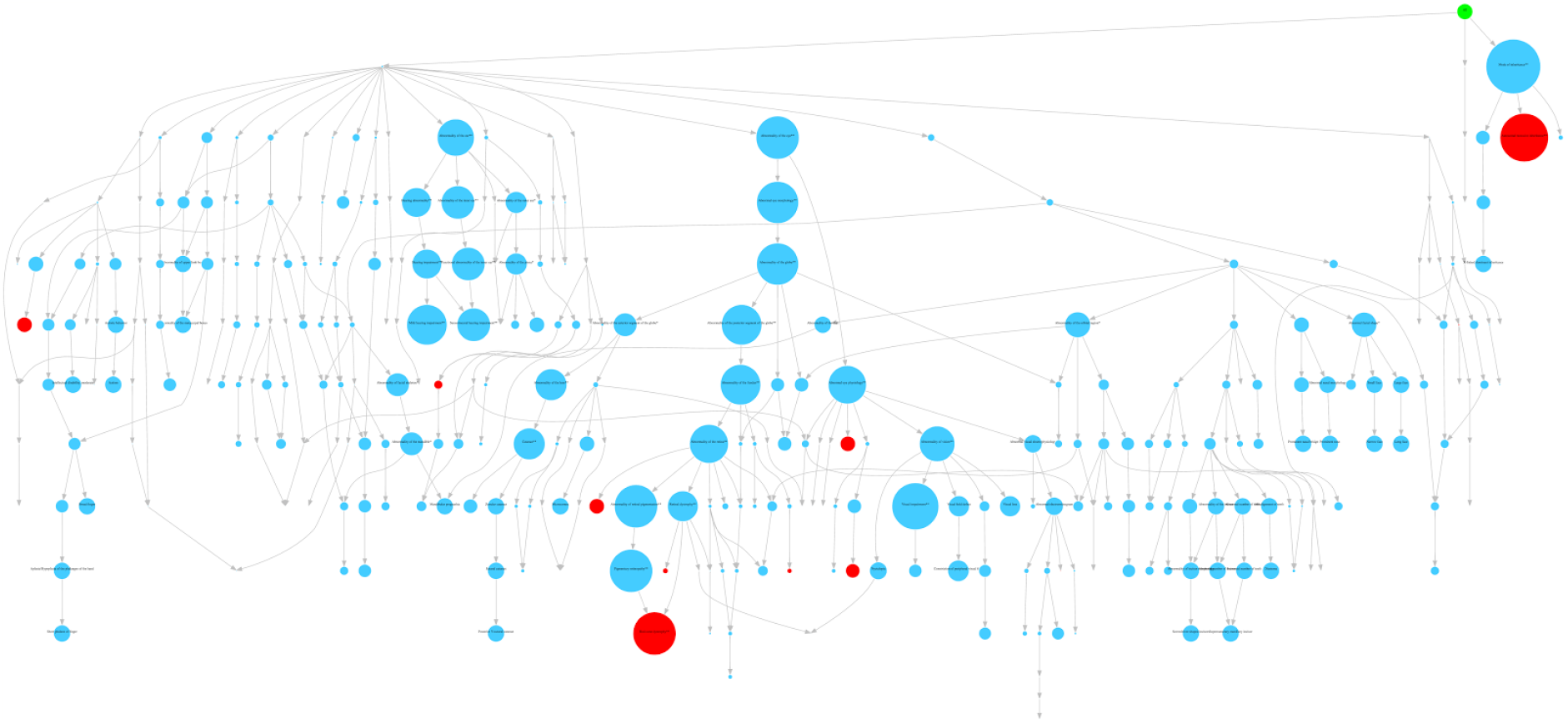

### SimReg

SimReg implements an inference procedure under the ‘similarity regression’ model described in (Greene et al. 2016). It is used here to estimate a probability of association between HPO encoded phenotypes and a binary genotype vector indicating whether a sequenced individual harbours a rare variant in a particular gene. This is done by comparing the evidence for a random model with one in which the log odds of observing a rare variant is linked to the similarity between HPO profiles and an estimated characteristic HPO profile. The method was applied to all genes under dominant and recessive mode of inheritance assumptions, i.e. setting the binary genotype to 1 if individuals carried at least 1 and 2 rare alleles respectively, and 0 otherwise. One application was to the gene *GUCY2D*, which was estimated to have a probability of association of 0.98. The estimated characteristic phenotype, shown below, corresponds well to the literature phenotype (Gregory-Evans et al. 2000).

SimReg is available for download on CRAN (https://cran.r-project.org/web/packages/SimReg/index.html).

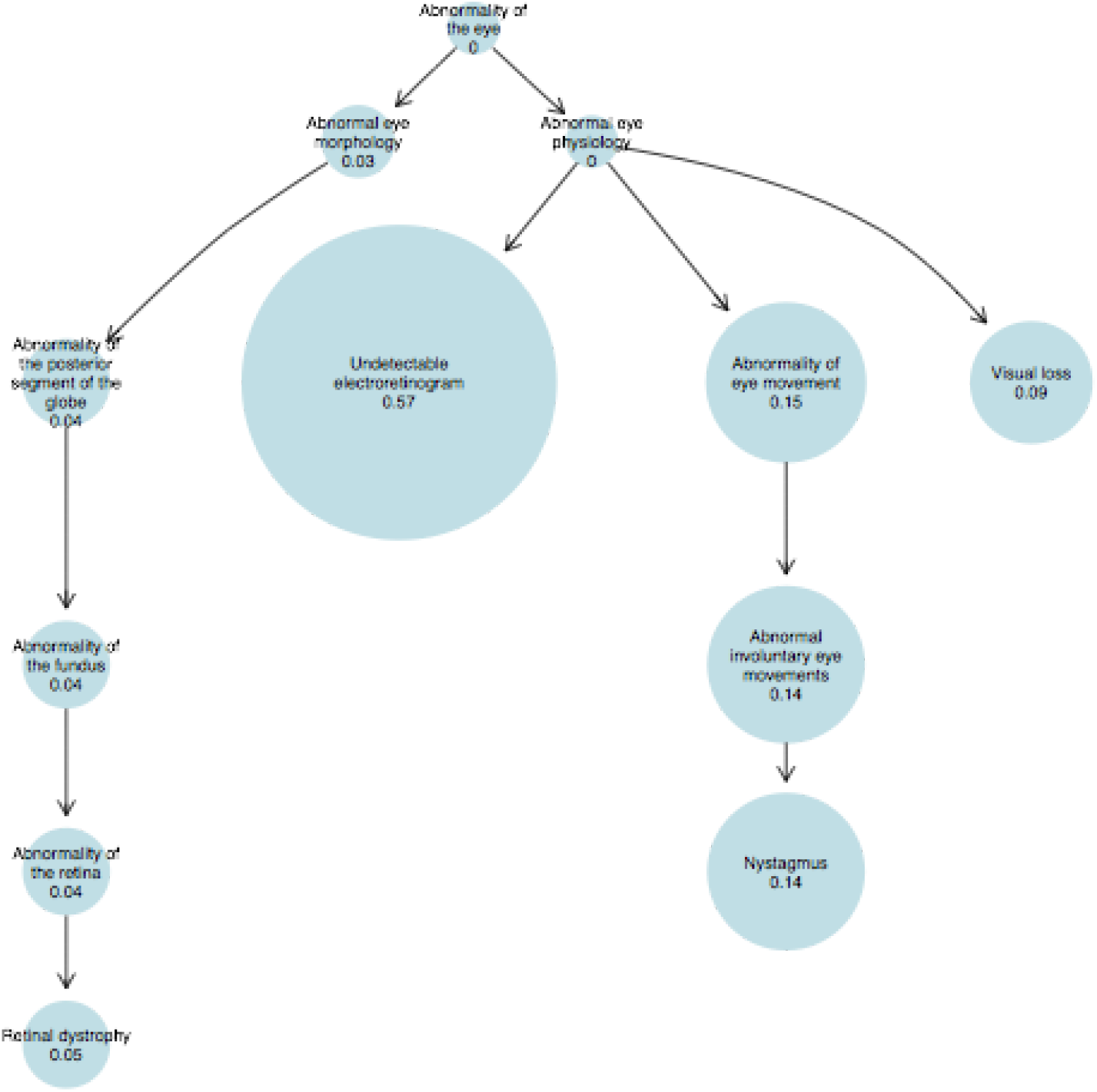

